# Faster growth enhances low carbon fuel and chemical production through gas fermentation

**DOI:** 10.1101/2022.02.13.480257

**Authors:** Lorena Azevedo de Lima, Henri Ingelman, Kush Brahmbhatt, Kristina Reinmets, Craig Barry, Audrey Harris, Esteban Marcellin, Michael Köpke, Kaspar Valgepea

## Abstract

Gas fermentation offers both fossil carbon-free sustainable production of fuels and chemicals and recycling of gaseous and solid waste using gas-fermenting microbes. Bioprocess development, systems-level analysis of biocatalyst metabolism, and engineering of cell factories are advancing the widespread deployment of the commercialised technology. Acetogens are particularly attractive biocatalysts but effects of the key physiological parameter – specific growth rate (μ) – on acetogen metabolism and the gas fermentation bioprocess have not been established yet. Here, we investigate the μ-dependent bioprocess performance of the model-acetogen *Clostridium autoethanogenum* in CO and syngas (CO+CO_2_+H_2_) grown chemostat cultures and assess systems-level metabolic responses using gas analysis, metabolomics, transcriptomics, and metabolic modelling. We were able to obtain steady-states up to μ ~2.8 day^-1^ (~0.12 h^-1^) and show that faster growth supports both higher yields and productivities for reduced by-products ethanol and 2,3-butanediol. Transcriptomics data revealed differential expression of 1,337 genes with increasing μ and suggest that *C. autoethanogenum* uses transcriptional regulation to a large extent for facilitating faster growth. Metabolic modelling showed significantly increased fluxes for faster growing cells that were, however, not accompanied by gene expression changes in key catabolic pathways for CO and H_2_ metabolism. Cells thus seem to maintain sufficient “baseline” gene expression to rapidly respond to CO and H_2_ availability without delays to kick-start metabolism. Our work advances understanding of transcriptional regulation in acetogens and shows that faster growth of the biocatalyst improves the gas fermentation bioprocess.

## INTRODUCTION

Climate change is causing alarming detrimental degradation to the environment. The world thus needs to decarbonize energy production (e.g. solar, wind) and move away from producing fuels and chemicals from fossil carbon. Furthermore, improved recycling of solid waste (e.g. municipal solid waste [MSW], plastic waste) is becoming increasingly important to maintain biosustainability. Gas fermentation offers a sustainable route for the production of renewable chemicals and fuels by recycling gaseous one-carbon (C1) waste feedstocks using gas-fermenting organisms (e.g. industrial waste gases, syngas from gasified MSW or biomass [CO+H_2_+CO_2_]) (Liew et al., 2016; Redl et al., 2017; Fackler et al., 2021; Pavan et al., 2022). Acetogen bacteria are particularly attractive for gas fermentation as they accept gas as their sole energy and carbon source (Wood 1991) and use the most efficient pathway to fix CO_2_ (Drake 2006; Fast and Papoutsakis 2012; Cotton et al., 2018), the Wood-Ljungdahl pathway (WLP) (Wood 1991; Ragsdale and Pierce, 2008). Acetogens generally convert carbon into acetic acid, ethanol, or 2,3-butanediol while metabolic engineering for expanding their product spectrum is advancing rapidly (Bourgade et al., 2021; Köpke and Simpson 2020; Fackler et al., 2021; Pavan et al., 2022). Notably, the acetogen *Clostridium autoethanogenum* is used as a commercial-scale gas fermentation cell factory (Köpke and Simpson, 2020).

A better understanding of acetogen metabolism and the gas fermentation bioprocess can contribute to the widespread deployment of the technology. Recent systems-level studies have improved the much-needed quantitative understanding of the energy-limited metabolism of acetogens (Schuchmann and Müller, 2014; Molitor et al., 2017) to advance their rational metabolic engineering. For example, regulatory principles behind metabolic shifts in carbon, energy, and redox balances (Valgepea et al., 2017a, 2018; Mahamkali et al., 2020; Richter et al., 2016), metabolic robustness (Mahamkali et al., 2020), and transcriptional and translational regulation (Lemgruber et al., 2019; Song et al., 2017; Al-Bassam et al., 2018; Song et al., 2018, Shin et al., 2021) have been quantified. At the same time, understanding of the gas fermentation bioprocess has also improved with the characterisation of the effects of gas-liquid mass transfer, feed gas and media composition, pH shifts, and mixed cultures on the biocatalyst and fermentation performance, e.g. gas uptake, biomass level, product profile, yields, and rates (Cotter et al., 2009; Abubackar et al., 2012; Abubackar et al., 2015; Valgepea et al., 2017b, 2018; Park et al., 2019; Heffernan et al., 2020; Esquivel-Elizondo et al., 2017; Diender et al., 2019). However, the effects of the specific growth rate (μ) of the cells on acetogen metabolism and on the gas fermentation bioprocess have not yet been established.

Characterisation of μ-dependent acetogen growth is important for three reasons. Firstly, it could reveal important insights into the energy-limited metabolism of acetogens (Schuchmann and Müller, 2014; Molitor et al., 2017) as faster growth demands more energy. Secondly, studies of the impact of μ on cell metabolism in other microorganisms have, for instance, revealed profound effects on product distribution and energy balance (Valgepea et al., 2010; Van Hoek et al., 1998), transcript and protein expression (Ishii et al., 2007; Valgepea et al., 2010, 2013; Peebo et al., 2015; Hackett et al., 2016), and stress responses (Regenberg et al., 2006). Thirdly, μ of the cell culture is an important bioprocess parameter affecting metabolic activity of cells, process rates, and economics (Lipson 2015). This is especially relevant for acetogen gas fermentation as the process is operated as a continuous culture at industrial-scale (Köpke and Simpson, 2020; Fackler et al., 2021). Thus, the selection of culture dilution rate (i.e. μ at steady-state) is critical for optimal bioprocess performance (e.g. titre, rate, yield).

In this work, we investigate the μ-dependent bioprocess performance of the acetogen *C. autoethanogenum* in CO- and syngas-grown chemostat cultures and assess systems-level metabolic responses using gas analysis, metabolomics, transcriptomics, and metabolic modelling. We obtained steady-states up to μ ~2.8 day^-1^ (~0.12 h^-1^) and show that faster growth supports both higher yields and productivities for reduced by-products ethanol and 2,3-butanediol (2,3-BDO), thereby benefitting the gas fermentation bioprocess. Transcriptomics data suggest that *C. autoethanogenum* uses transcriptional regulation to a large extent for facilitating faster growth and emphasise the need for mapping genotype-phenotype links and improving gene annotations in acetogens for advancing understanding of metabolism and engineering of cell factories.

## MATERIAL AND METHODS

### Bacterial strain, growth medium, and continuous culture conditions

A derivate of *Clostridium autoethanogenum* DSM 10061 strain – DSM 23693 – deposited in the German Collection of Microorganisms and Cell Cultures (DSMZ) was used in all experiments and stored as a glycerol stock at −80°C. Cells were grown either on CO (60% CO and 40% Ar; AS Eesti AGA) or syngas (50% CO, 20% H_2_, 20% CO_2_, and 10% Ar; AS Eesti AGA) in a chemically defined medium without yeast extract described before (Valgepea et al., 2017a). Cells were grown under strictly anaerobic conditions at 37°C and at pH 5 maintained by 5M NH_4_OH. Chemostat continuous cultures were performed in 1.4 L Multifors bioreactors (Infors AG) at a working volume of 750 mL connected to a Hiden HPR-20-QIC mass spectrometer (Hiden Analytical) for online high-resolution off-gas analysis. The system was equipped with peristaltic pumps; mass flow controllers (MFCs); pH, ORP, and temperature sensors. Antifoam (435530; Sigma-Aldrich) was continuously added to the bioreactor at a rate of 10 μL/h to avoid foaming. In total, 23 chemostat cultures were run at three dilution rates: ~1.0, ~2.0, and ~2.8 day^-1^ (μ~0.04, 0.08, and 0.12 h^-1^, respectively) with variable gas-liquid mass transfer rates to maintain similar steady-state biomass concentrations (**Table 1**). All steady-state results reported here were collected after optical density (OD), gas uptake, and production rates had been stable for at least 3–5 working volumes.

**Table 1.** Variable operational parameters of 23 steady-state chemostat cultures of *C. autoethanogenum*.

| Feed gas | Gas flow rate (mL/min) | Agitation (RPM) | $\mu$ (day<sup>-1</sup>) | Biomass concentration (gDCW/L) | Number of bioreplicates |
| --- | --- | --- | --- | --- | --- |
| CO | 50 | 690 | 1.02 ± 0.01 | 1.58 ± 0.06 | 4 |
|  | 72 | 815 | 2.03 ± 0.06 | 1.65 ± 0.01 | 3 |
|  | 72 | 1175 | 2.79 ± 0.03 | 1.65 ± 0.02 | 3 |
| Syngas | 50 | 675 | 1.01 ± 0.02 | 1.59 ± 0.03 | 6 |
|  | 72 | 800 | 2.01 ± 0.07 | 1.57 ± 0.08 | 3 |
|  | 72 | 1160 | 2.79 ± 0.07 | 1.43 ± 0.03 | 4 |

$\mu$ , specific growth rate; gDCW, gram of dry cell weight. Data are average ± standard deviation between bioreplicates.

### Biomass concentration Analysis

Biomass concentration in gram of dry cell weight per litre of broth (gDCW/L) was determined by measuring the OD of the culture at 600 nm after the correlation coefficient (K) between culture OD and DCW was established at 0.23, using the methodology described in Peebo et al., 2014.

### Extracellular metabolome analysis

Analysis of exo-metabolome was performed using filtered broth samples stored at −20°C until analysis. Organic acids and alcohols were analysed by HPLC (Shimadzu Prominence-I LC-2030 plus system) using a Rezex™ ROA-Organic Acids H^+^ (8%) 300 × 7.8 mm column (00H-0138-K0; Phenomenex) and a guard column (03B-0138-K0; Phenomenex). Twenty microlitres of the sample were injected using an auto-sampler and eluted isocratically with 0.5 mM H_2_SO4 at 0.6 mL/min for 30 min at 45°C. Compounds were detected by a refractive index detector (RID-20A; Shimadzu) and identified and quantified using relevant standards using the software LabSolution (Shimadzu). We note that cells produced 2R,3R-butanediol.

### Bioreactor off-gas analysis

Bioreactor off-gas analysis was performed by an online Hiden HPR-20-QIC mass spectrometer and specific gas uptake (CO and H_2_) and production rates (CO_2_, ethanol) (mmol/gDCW/h) were determined as described before (Valgepea et al., 2017a). The Faraday Cup detector monitored the intensities of H_2_, CO, ethanol, H_2_S, Ar, and CO_2_ at 2, 14, 31, 34, 40, and 44 amu, respectively.

### Carbon balance analysis

Carbon recoveries and balances were determined as described before (Valgepea et al., 2017a).

### Transcriptome analysis

Transcriptome analysis of 20 chemostat cultures was conducted using RNA sequencing (two and one bio-replicate of syngas cultures at ~1 and ~2.8 day^-1^, respectively, indicated in **Table 1** were excluded due to experimental constraints). Ten millilitres of culture were pelleted by immediate centrifugation (5,000 × *g* for 3 min at 4°C) and resuspended in 5 mL of RNAlater (76106; Qiagen). Samples were stored at 4°C overnight, centrifuged (4,000 × *g* for 10min at 4°C), and pellets stored at −80°C until RNA extraction.

Thawed cell pellets were resuspended in 800 μl of RLT buffer (74104; Qiagen) containing 10 μl of β-mercaptoethanol and lysed with acid-washed glass beads (G4649; Merck) using the Precellys^®^ 24 instrument with liquid nitrogen cooling (Bertin Technologies). Total RNA was extracted using the RNeasy mini kit (74104, Qiagen) with off-column TURBO1™ DNase treatment (AM2239; Invitrogen), followed by purification and enrichment using the RNA Clean and Concentrator™ kit (R1018, Zymo). The efficiency of the total RNA purification and DNA removal was verified using the NanoDrop™ 1000 instrument (Thermo Scientific) and the quality of RNA extracts was checked using the TapeStation 2200 equipment (Agilent Technologies). Total RNA concentration was determined using the Qubit 2.0 instrument (Q32866; Invitrogen). Next, ribosomal RNA (rRNA) was removed using the QIAseq FastSelect −5S/16S/23S Kit (335925; Qiagen) and stranded mRNA libraries were prepared using the QIAseq Stranded RNA Lib Kit (180743; Qiagen). RNA sequencing was performed using the NextSeq MID150 sequencing kit (20024904; Illumina) on the NextSeq500 sequencer (Ilumina) with 2 x 75 bp paired-end dual indexed (2 x 8 bp) reads, which produced eight fastq files per sample (160 files in total).

### RNA sequencing data analysis

The R-scripts used for the analysis of RNA sequencing data after read trimming includes complete details of the methodology and can be downloaded as **Supplementary Files 1** and **2**.

#### Mapping and assignment of genome features from RNA sequencing raw data

The quality of raw NextSeq reads was verified using MultiQC (Ewels et al., 2016) and adapter sequences were trimmed using the Cutadapt Python package (version 2.10; Martin 2011) allowing a minimum read length of 35 nucleotides. The resulting high-quality paired-end reads were mapped to the NCBI reference genome NC_022592.1 (Brown et al., 2014) using the align function within Rsubread package (version 2.4.2; Liao et al., 2019). Thereafter, four .bam files per sample were merged using Samtools (version 1.10; Li et al., 2009) and genomic features were assigned using the featureCounts functions within Rsubread. Samples had 5.6–9.3 million reads with an average mapping rate of 99% and this generated 4.3–6.4 million feature counts across samples (**Table S1**). The NCBI annotation NC_022592.1 of the *C. autoethanogenum* sequence (Brown et al., 2014) was used as the annotation genome, including only coding (CDS) and non-coding (ncRNA) sequences. Additionally, CAETHG_RS07860 was removed from the annotation and replaced with the carbon monoxide dehydrogenase genes with initial IDs of CAETHG_1620 and 1621 (Brown et al., 2014) which were given the IDs CAETHG_RS07861 and RS07862, respectively.

#### Determination of transcript abundances and differentially expressed genes (DEGs)

Transcript abundances and DEGs were determined as described before (Valgepea et al., 2017a). In short, raw library sizes were normalized and transcript abundances in reads per kilobase of transcript per million mapped reads (RPKM) were estimated from feature counts and gene lengths using edgeR (version 3.32.1; Robison et al., 2010). Transcripts with abundances >10 RPKMs in at least two samples were subject to differential expression analysis using limma (version 3.46.0; Ritchie et al., 2015) between bio-replicate cultures of different μ values within one gas mixture (i.e. gas mixes at same μ were not compared). DEGs were determined by fold-change >1.5 and q-value < 0.05 after false discovery rate (FDR) correction (Benjamini and Hochberg, 1995). Transcript abundances and DEGs are presented in **Tables S2**, **S3** and **S4**, **S5**, respectively. Proposed gene names were obtained from Valgepea et al. 2021. RNA sequencing data (NextSeq) has been deposited in the NCBI Gene Expression Omnibus repository under accession number GSEXXX.

#### Functional data analysis

Mapping of gene IDs to Cluster of Orthologous Groups (COGs) was performed using the eggNOG 5.0 database (Huerta-Cepas et al., 2019) that resulted in COG assignment for 1,190 genes (**Table S6**). The Gene Ontology (GO) terms list was assembled using Pannzer2 (Törönen et al., 2018), InterProScan5 (Jones et al., 2014), and eggNOG 5.0 databases that resulted in GO term assignment for 3,001 genes (**Table S7**). Clustering of transcript expression profiles across μ values and gas mixes in **Figure 5B** was performed and visualised using the dendextend (version 1.14.0; Galili 2015) and ComplexHeatmap (version 2.10.0; Gu et al., 2016) packages in R using Ward’s ward.D2 hierarchical clustering algorithm. Six clusters were identified based on the sharp decline of the dendrogram height parameter for other clusters. Functional enrichment analysis of GO terms was performed using the topGO package in R (version 2.40.0; Alexa and Rahnenfuhrer, 2020) and a custom-made script (**Supplementary File 2**) with significant enrichment indicated by q-Fisher < 0.05 for FDR-corrected Fisher’s exact test. The 245 genes that quantitatively showed the same expression trend with increasing μ on both gas mixes (**Figure 5C**) were determined by a t-test (p-value < 0.05) between the slopes of linear regression of gene expression change on the two gas mixes for genes that showed a continuously increasing or decreasing differential expression (i.e. DEGs for both μ = 2.0 vs. 1.0 and 2.8 vs. 2.0 day^-1^) with increasing μ on both gases.

### Genome-scale metabolic modelling

Model simulations were performed using the genome-scale model iCLAU786 of *C. autoethanogenum* (Valgepea et al., 2018) and flux balance analysis (FBA) (Orth et al., 2010) as specified in Valgepea et al. (2018). Briefly, intracellular metabolic flux rates (SIM1–23 in **Tables S8, S9**) were estimated using experimentally measured constraints (e.g. μ, substrate consumption and product production rates) and maximisation of ATP dissipation as the objective function in FBA. Prediction of optimal growth phenotypes for μ and products (SIM24–46) was performed using experimental substrate uptake rates, the ATP dissipation flux calculated above, and maximisation of biomass yield as the objective function. Accuracy of growth phenotype prediction was improved (SIM47–59) by additionally zeroing CO_2_ reduction with the redox-consuming FdhA activity (reaction rxn00103_c0) and fixing the ratio between H_2_ utilisation for direct CO_2_ reduction (reaction rxn08518_c0), and Fd_red_ and NADPH generation (reaction leq000001) by the HytA-E/FdhA complex at a value corresponding to the respective experiment’s q_H2_/q_CO_ ratio (see Valgepea et al., 2018 for details). Simulation results identified as SIMx (e.g., SIM1) in the text are reported in **Tables S8** and **S9**, while the SBML model file of iCLAU786 is supplied in Valgepea et al. (2018).

## RESULTS

### Steady-state gas-fermenting chemostat cultures of *Clostridium autoethanogenum*

We employed continuous cultures for controlling the μ of cells as this allows to unequivocally quantify the effects of μ on cell growth compared to batch cultures where genetically modified strains (e.g. titratable substrate uptake) or variable culture parameters (e.g. growth media) need to be used that might confound results (Adamberg et al., 2015). Here, the acetogen *Clostridium autoethanogenum* was grown in 23 autotrophic chemostats where cells reached steady-states on CO or syngas (CO+CO_2_+H_2_) at 37 °C and pH 5 using a chemically defined medium in biological triplicates or quadruplicates at dilution rates ~1.0, ~2.0, and ~2.8 day^-1^ (μ~0.04, 0.08, and 0.12 h^-1^, respectively) (**Table 1**). In addition to continuous quantification of gas uptake and production rates using an online mass spectrometer, cultures were sampled for extracellular metabolome and transcriptome analysis. We also used genome-scale metabolic modelling to estimate intracellular flux rates.

### Elevated ethanol productivity with faster growth

We adjusted gas-liquid mass transfer between different dilution rates to maintain similar steady-state biomass concentrations (~1.6 ± 0.1 gDCW/L; average ± standard deviation) (**Table 1** and **Figure 1A**) as it can affect acetogen carbon distribution (Valgepea et al., 2017a). We detected secretion of acetate, ethanol, and 2,3-BDO by the cells (**Figure 1A**). While specific acetate production rate (q_ace_; mmol/gDCW/h) profiles with increasing μ varied between the two gases, specific ethanol production rates (q_EtOH_) more than doubled on both gases (**Figure 1B**). This also means higher productivity (g/L/h) due to similar biomass levels. Increasing trends for specific 2,3-BDO production rates (q_2,3-BDO_) were also seen with higher μ. Notably, faster growth led to molar acetate/ethanol ratios dropping from ~1.5 to 0.9 and 0.6 for CO and syngas, respectively, due to a strong decrease of acetate levels at similar ethanol values (**Figure 1A**). These trends are beneficial for an industrial ethanol production process as higher productivity is not comprised with lower selectivity.

**Figure 1.**
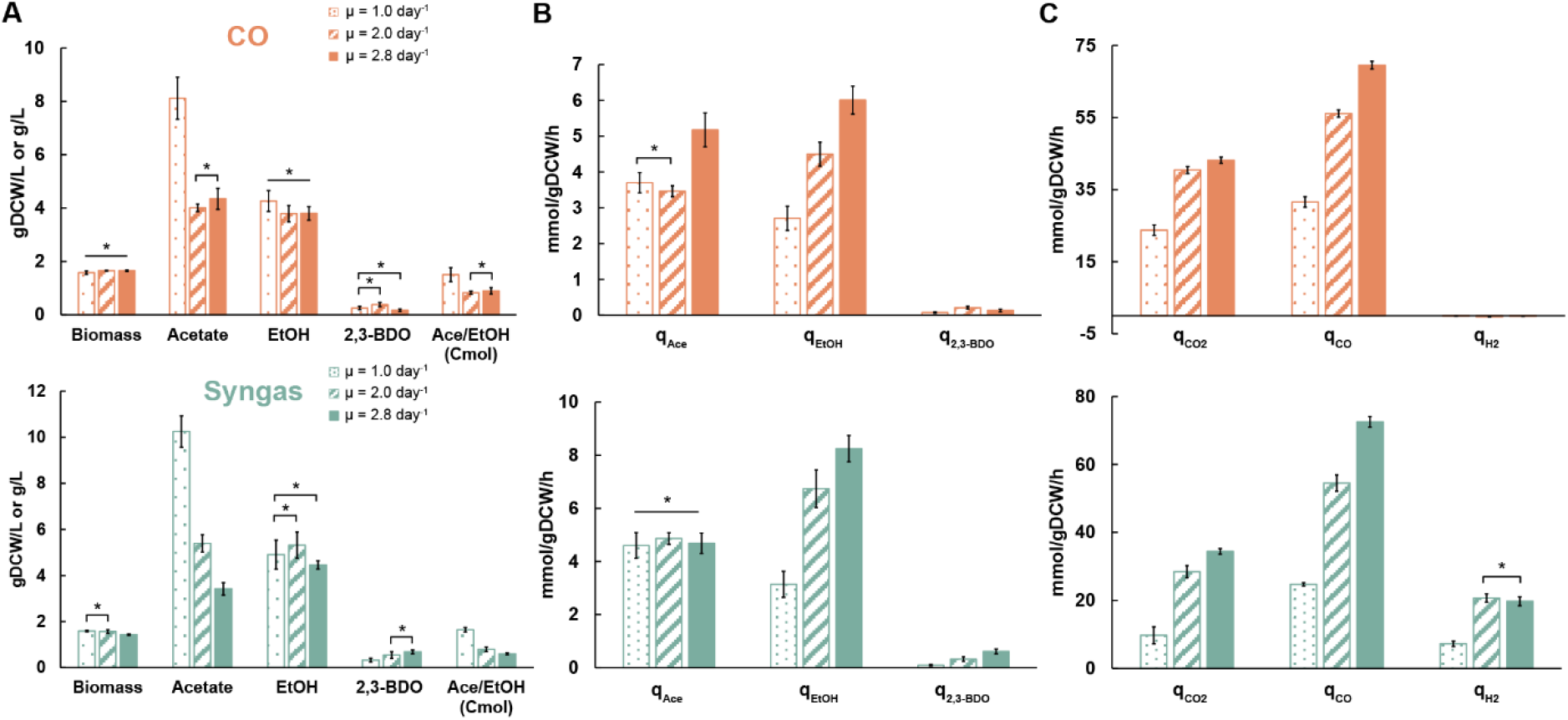
Specific growth rate-dependent growth characteristics in CO (top) and syngas (bottom) grown *C. autoethanogenum* chemostats. **(A)** Biomass and by-product concentrations. **(B)** Specific by-product production rates. **(C)** Specific gas uptake and production rates. Data are average ± standard deviation between bio-replicates (see **Table 1**). Asterisk denotes values statistically not different according to t-test (p-value < 0.05). *μ*, specific growth rate; *gDCW*, gram of dry cell weight; *EtOH*, ethanol; *2,3-BDO*, 2,3-butanediol; *Ace*, acetate; *q*, specific rate; *qCO2*, specific CO_2_ production rate; *qH2* and *qH2*, specific CO and H_2_ uptake rates, respectively.

### Gas analysis indicates metabolic rearrangements

CO limited our chemostats since we could maintain similar steady-state biomass concentrations across dilution rates by changing gas-liquid mass transfer (**Table 1**). One could thus expect the specific CO uptake rate (qCO; mmol/gDCW/h) to increase proportionally with μ. We detected qCO to increase ~2.2-(32 ± 1 to 70 ± 1) and ~2.9-fold (25 ± 1 to 73 ± 2) for CO and syngas cultures, respectively, with 2.8-fold faster growth (**Figure 1C**). The non-proportional change for CO cultures means a higher biomass yield (gDCW/mmol of CO consumed) at higher μ. As expected for CO-limited chemostats (Richter et al., 2013; Martin et al., 2015; Valgepea et al., 2018), simultaneous uptake of CO and H_2_ were observed for syngas cultures (**Figure 1C**). Interestingly, while the specific H_2_ uptake rate (qH_2_; mmol/gDCW/h) increased ~2.9-fold (7 ± 1 to 21 ± 1) between μ = 1.0 and 2.0 day^-1^ on syngas, no further change was seen with faster growth, despite gas-liquid mass transfer being increased proportionally with dilution rate. Specific CO_2_ production rates (qCO2; mmol/gDCW/h) increased by ~1.8- (24 ± 1 to 43 ± 1) and ~3.4-fold (10 ± 2 to 34 ± 1) for the CO and syngas cultures, respectively (**Figure 1C**), indicating a strong effect of H_2_ uptake on CO oxidation as seen before for *C. autoethanogenum* growing on various gas mixes (Valgepea et al., 2018).

### Faster growth leads to carbon diversion away from acetate

Product ratios show the distribution of carbon between products but carbon balancing is necessary to quantify carbon flows from substrates to products (i.e. yields). Notably, carbon flux to acetate dropped with faster growth by 24% (19.2 ± 2.2% to 14.6 ± 1.3%) on CO and by 57% (32.3 ± 2.5% to 13.9 ± 1.1%) on syngas (**Figure 2**). While no clear μ-dependent trend for carbon flow to 2,3-BDO was detected on CO, cells diverted up to 141% more carbon into 2,3-BDO in syngas cultures (1.5 ± 0.3% to 3.6 ± 0.4%). The aforementioned doubling of q_EtOH_ with faster growth (**Figure 1B**) was supported by up to ~21% elevated carbon flux to ethanol (**Figure 2**). These observations are not trivial as faster growth demands more energy while reduced products like ethanol and 2,3-BDO consume redox co-factors that acetogens could otherwise use for ATP generation (Schuchmann and Müller, 2014). In fact, cells were even able to channel more carbon into biomass formation simultaneously (increase of ~48 and ~24% for CO and syngas, respectively; **Figure 2**). The ~60% and ~50% losses of substrate carbon as CO_2_ for the CO and syngas cultures, respectively, are close to theoretical stoichiometric and thermodynamic calculations for CO_2_ dissipation with ethanol production from CO (~67%) or from a gas mix with a CO-to-H_2_ ratio around two (50%) (Wilkins and Atiyeh, 2011; Molitor et al., 2016).

**Figure 2.**
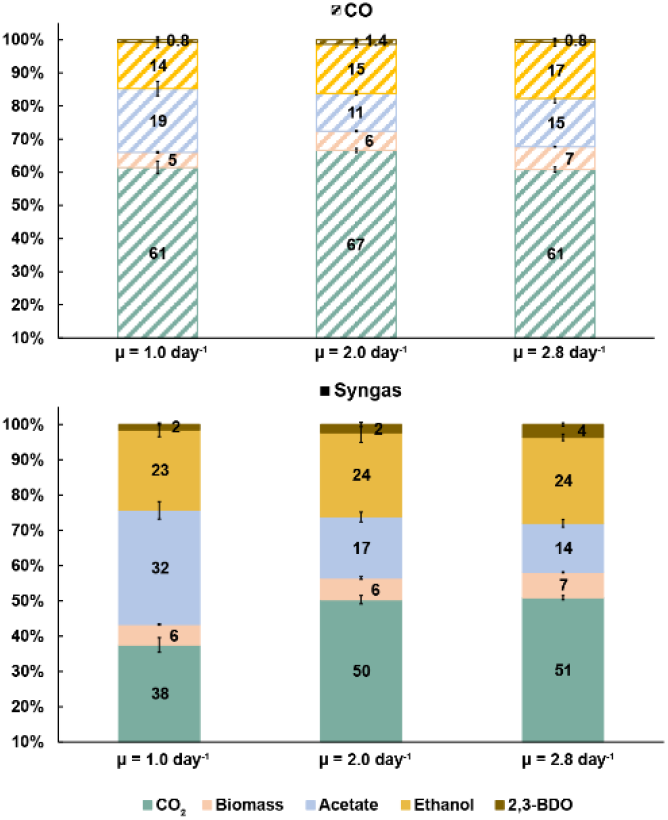
Specific growth rate-dependent carbon balances in CO (top) and syngas (bottom) grown *C. autoethanogenum* chemostats. Carbon recoveries were 110 ± 16%, 103 ± 6%, 92 ± 1% for CO and 122 ± 6%, 107 ± 3%, 101 ± 3% for syngas at μ = 1.0, 2.0, 2.8 day^-1^, respectively. Carbon recoveries were normalized to 100% to have a fair comparison of carbon distributions between different conditions. Data are average ± standard deviation between bioreplicates (see **Table 1**). *μ*, specific growth rate; *2,3-BDO*, 2,3-butanediol.

### Analysis of metabolic fluxes using a genome-scale metabolic model

We took advantage of the quantitative steady-state data collected above on carbon and redox flows entering and leaving the cells to estimate intracellular metabolic fluxes using the genome-scale metabolic model (GEM) iCLAU786 of *C. autoethanogenum* (Valgepea et al., 2018) and flux balance analysis (FBA) (Orth et al., 2010). Steady-state intracellular flux patterns for our CO and syngas chemostats were estimated by constraining the GEM with experimental data (exchange rates and μ) and maximising ATP dissipation as the objective function in FBA (SIM1–23 in **Tables S8, S9**).

Flux through the WLP was significantly increased with faster growth, although less than the increase in CO uptake for syngas cultures as more of the fixed CO was oxidised and dissipated as CO_2_ (**Figure 3** and **Tables S8, S9**). Faster growth was supported with elevated ATP production through the ATPase (by 82 and 125% on CO and syngas, respectively; **Figure 3**). Interestingly, the ratio between the ATP production fluxes of the ATPase and the acetate kinase (acetyl-P → acetate) changed minimally (9% in average; **Tables S8**), despite significantly altered product distributions (**Figure 2**). At the same time, maintenance ATP costs increased by 80 and 66% to 9.9 ± 0.5 and 11.7 ± 1.9 mmol/gDCW/h on CO and syngas, respectively (**Figure 3**). The average ~30% fraction of maintenance costs from total ATP production (**Table S8**) is similar to previously studied autotrophic *C. autoethanogenum* cultures (Valgepea et al., 2017a, 2018). In addition to ATP, biomass synthesis during faster growth also demands an elevated supply of NADPH. Extra NADPH was provided by increased flux through the Nfn transhydrogenase (by 66 and 190% to 12.0 ± 0.2 and 8.8 ± 0.5 mmol/gDCW/h on CO and syngas, respectively), and not by higher flux through the electron-bifurcating hydrogenase HytA-E complex on syngas (**Figure 3**). No increase in flux through HytA-E during faster growth on syngas suggested that production of the critical reducing power co-factor, reduced ferredoxin (Fdred), was possibly supported by CO oxidation. Indeed, the model demonstrated an increase of the fraction of Fdred generated by CO oxidation from 63 ± 2% to 74 ± 1% with increasing μ (**Table S8**). As seen for previous CO+H_2_ or CO_2_+H_2_-fermenting *C. autoethanogenum* steady-state cultures (Valgepea et al., 2017a, 2018; Heffernan et al., 2020), all the CO_2_ in the WLP was reduced to formate directly using H_2_ through the formate-H_2_ lyase activity of the HytA-E/formate dehydrogenase (FdhA) complex (Wang et al., 2013). This saves valuable redox compared to growth on CO only and allows higher production of reduced products when H_2_ is available (**Figure 2**). Regarding ethanol production, our simulations are consistent with the aforementioned *C. autoethanogenum* datasets and gene knockout experiments (Liew et al., 2017) showing that ethanol was synthesised using the aldehyde:Fd oxidoreductase (AOR) activity to couple it with ATP production (**Figure 3**).

**Figure 3.**
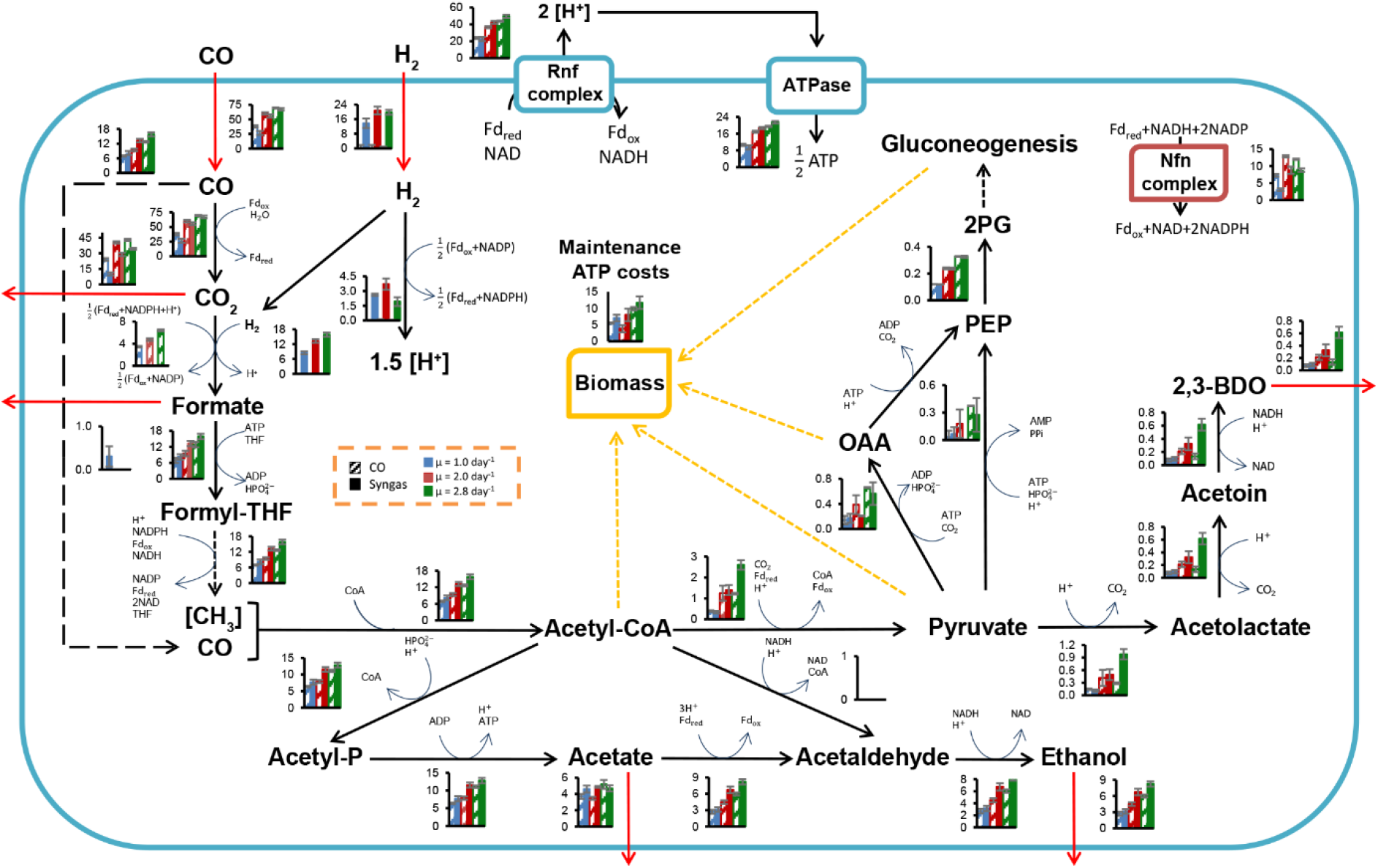
Specific growth rate-dependent central metabolism flux levels in CO and syngas grown *C. autoethanogenum* chemostats. See dashed inset for bar chart details. Fluxes (mmol/gDCW/h) are represented as average ± standard deviation between bioreplicates (see **Table 1**). Arrows show direction of calculated fluxes; red arrows denote uptake or secretion; dashed arrows denote a series of reactions. Cofactors used in the GEM iCLAU786 are shown. Flux into PEP from oxaloacetate and pyruvate is merged. *gDCW*, gram of dry cell weight; *μ*, specific growth rate; *OAA*, oxaloacetate; *PEP*, phosphoenolpyruvate; *THF*, tetrahydrofolate; *2PG*, 2-phosphoglycerate; *2,3-BDO*, 2,3-butanediol. See SIM1–23 in **Tables S8, S9** for data and co-factor abbreviations.

GEMs can also be used for predicting growth phenotypes (O’Brien et al., 2015) and the accuracy of the model can be evaluated beforehand, for example, by comparing experimental product profiles with predicted ones if constraining only substrate uptake rates, maintenance ATP costs, and maximising biomass yield in FBA. We initially failed to predict ethanol and 2,3-BDO production (SIM24–46 in **Tables S8, S9**), as seen before (Valgepea et al., 2017a, 2018). However, when additionally constraining the model by coupling carbon and redox metabolism from H_2_ utilisation for syngas cultures (see Materials and Methods), ethanol production was predicted on average 28% off from experimental values (SIM47–59 in **Tables S8, S9**). Prediction of acetogen phenotypes can be further improved by considering thermodynamics and kinetics (Greene et al., 2019; Mahamkali et al., 2020), which are important for accounting for the effects from high extracellular product levels (e.g. acetate, ethanol).

### Global transcriptome trends with faster growth

Next, we performed transcriptome analysis using RNA sequencing (RNA-seq) to quantify μ-dependent gene expression changes on both gas mixes. High reproducibility of the data was demonstrated by clear clustering of bio-replicates (**Figures 4A, B**) and an average Pearson correlation coefficient of R = 0.96 between bio-replicates across μ values (**Figure 4C**). Also, significant μ-dependent expression differences for many genes could be seen for both gases (**Figure 4B**). Indeed, we determined 1,337 differentially expressed genes (DEGs) in total with a fold-change (FC) > 1.5 (up- or down-regulation) and a q-value < 0.05 after FDR correction in at least one pairwise comparison within the three pairwise μ-dependent comparisons (μ = 2.0 vs. 1.0, 2.8 vs. 1.0, and 2.8 vs. 2.0 day^-1^) of both gas mixes (**Tables S4, S5**). Expectedly, the bigger the difference between compared μ values, the higher the number of DEGs (**Figure 4D**). Also, many DEGs were shared between comparisons within both gas mixes. Interestingly, while 442 DEGs were shared between CO and syngas comparisons of μ = 2.8 vs. 1.0 day^−1^, hundreds of unique DEGs were also detected, demonstrating the effect of the feed gas mix on μ-dependent gene expression patterns.

**Figure 4.**
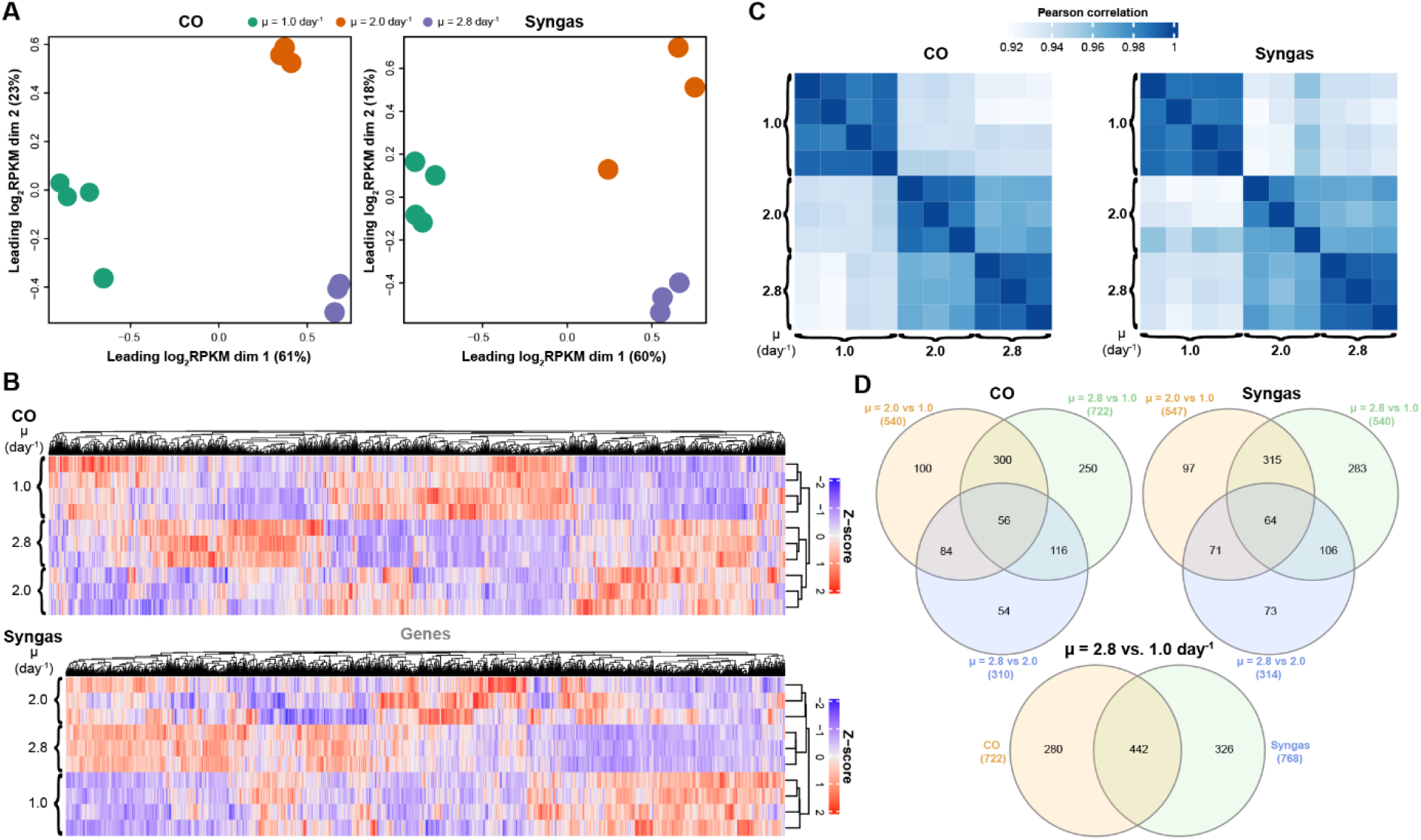
Specific growth rate-dependent transcriptome characteristics in CO- and syngas-grown *C. autoethanogenum* chemostats. **(A)** Multidimensional scaling (MDS) of bioreplicate transcript abundances (RPKM). **(B)** Hierarchical clustering of bioreplicate transcript abundances (Z-scores based on RPKMs). **(C)** Correlation of bioreplicate transcript abundances (RPKM) **(D)** Venn diagrams showing overlap of DEGs (number of DEGs for each comparison in brackets). *RPKM*, reads per kilobase of transcript per million mapped reads; *DEG*, differentially expressed gene (fold-change > 1.5 with q-value < 0.05); *μ*, specific growth rate. See **Tables S2, S3** and **S4, S5** for RPKM and DEG data, respectively.

**Figure 5.**
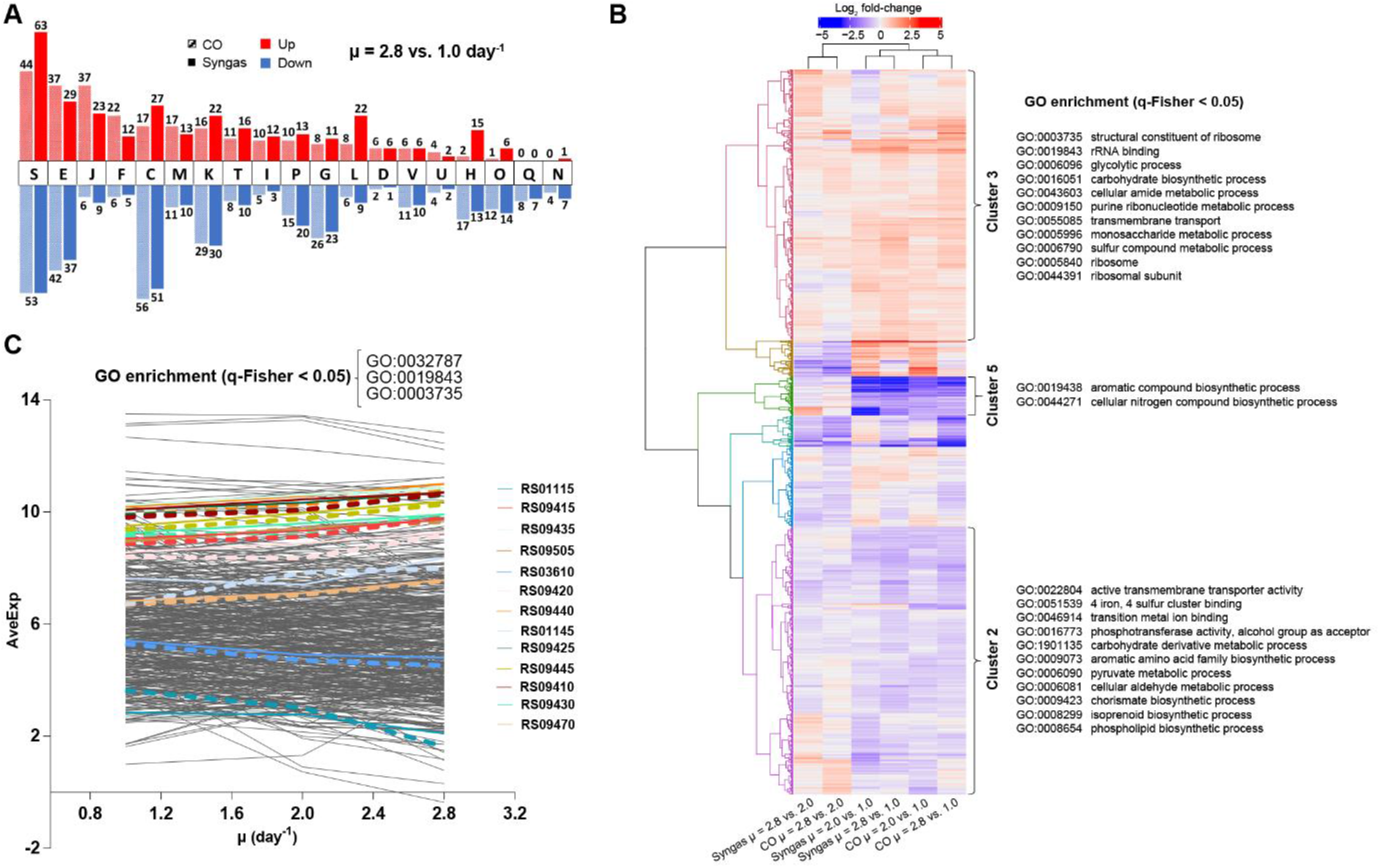
Global transcriptome changes assessed through functional gene classifications in CO- and syngas-grown *C. autoethanogenum* chemostats. **(A)** COG classification of DEGs for μ = 2.8 vs. 1.0 day^-1^ (see **Table S6** for COG descriptions). **(B)** Hierarchical clustering and GO enrichment of expression changes of all 1,337 DEGs. Six identified expression clusters are illustrated with the coloured dendrogram with three clusters showing GO enrichment (clusters 2, 3, and 5). **(C)** Genes and GO terms with tightly controlled expression with faster growth. All 245 genes showing the same quantitative expression trend on both gas mixes are shown with 13 genes within the enriched GO terms denoted by coloured lines (CO, dashed; syngas, solid). GO:0032787, monocarboxylic acid metabolic process. See **(B)** for other GO terms. Gene IDs are preceded with CAETHG_. *COG*, Cluster of Orthologous Groups; *DEG*, differentially expressed gene (fold-change > 1.5 with q-value < 0.05); *μ*, specific growth rate; *GO*, Gene Ontology; *q-Fisher*, FDR-corrected p-value of Fisher’s exact test; *AveExp*, average of bioreplicates log_2_ counts per million mapped reads (CPM; see Ritchie et al., 2015). See **Tables S4, S5** and **S7** for DEG data and GO terms list, respectively.

To understand global transcriptome changes that facilitate faster growth, we assessed DEGs through functional gene classifications. Firstly, analysis of DEGs of μ = 2.8 vs. 1.0 day^-1^ using Cluster of Orthologous Groups (COG) classification (Tatusov et al., 2003) showed that most DEGs were classified as “Function unknown” (S) (**Figure 5A** and **Table S6**). This highlights the need for genotype-phenotype mapping and improving gene annotations for acetogens. Genes involved in energy, amino acid, and nucleotide metabolism (C, E, and F) and replication, transcription, and translation (J, K, and L) were also abundant among DEGs and all these processes are vital for faster growth. Secondly, we used Gene Ontology (GO) terms (Ashburner et al., 2000) to attain finer resolution behind global gene expression trends. Clustering of all 1,337 DEGs across μ comparisons and gas mixes resulted in six significant expression clusters (**Figure 5B**). GO enrichment analysis (q-Fisher < 0.05) of the genes within the identified clusters was consistent with COG analysis as several GO terms related to translation (0003735, 0019843, 0005840, and 0044391) and metabolism (0006096, 0016051, 0043603, 0009150, 00505996 and 0006790) were enriched in cluster 3 that includes up-regulated genes with faster growth. Enrichment of GO terms within clusters 2 and 5 was detected, including genes down-regulated with faster growth. Lastly, we aimed to identify genes and GO terms whose expression was tightly controlled with changing μ, indicated by the same expression trend on both gas mixes. We identified 245 such genes that showed either continuously increasing or decreasing differential expression (DEGs, *i.e*. FC > 1.5 and q < 0.05) with increasing μ on both gases with the same magnitude of expression change between the two gas mixes (**Figure 5C**). Enrichment analysis of these genes confirmed the aforementioned results by identifying enrichment of genes (q-Fisher < 0.05) associated with translation (0003735 and 0019843), but also of the GO term 0032787 “monocarboxylic acid metabolic process” (coloured lines on **Figure 5C**).

### Transcriptional changes linked to metabolic rearrangements

Previous transcriptomics and proteomics studies have suggested that autotrophic metabolism of acetogens is not controlled through hierarchical regulation of gene expression (Richter et al., 2015; Valgepea et al., 2017a, 2018; Al-Bassam et al., 2018; Mahamkali et al., 2020). We thus analysed DEGs in terms of individual genes linked to activities specifically relevant to acetogens or with high expression changes. Surprisingly, we found numerous DEGs in central metabolism, some of which can explain the metabolic rearrangements described above (**Figures 6, S1, S2**). For instance, increased carbon flux to 2,3-BDO with faster growth on syngas (**Figure 2**) was supported by 18.5-fold (q < 0.01) (all μ = 2.8 vs. 1.0 day^-1^ unless otherwise noted) up-regulation of the 2,3-BDO dehydrogenase (CAETHG_RS01830; BDH) that reduces acetoin to 2,3-BDO and by 2.3-fold (q < 0.01) up-regulation of the pyruvate:Fd (flavodoxin) oxidoreductase (RS14890; PFOR) that converts acetyl-CoA to pyruvate (**Figure 6**). At the same time, elevated production of ethanol (**Figure 2**) could be linked to increased expression (2.1- to 2.8-fold, q < 0.01) of several alcohol dehydrogenases (RS02620 and RS02630; Adh and Adh3), including the most abundant – RS08920; Adh4 – in *C. autoethanogenum* (**Table S4**) (Valgepea et al., 2017a, 2021) on syngas (**Figure 6**). Acetaldehyde for the alcohol dehydrogenases was likely not supplied directly from acetyl-CoA as we quantified strong repression of two putative acetaldehyde dehydrogenases (RS08810 and RS08865; 11.5- and 17.0-fold, q < 0.01), which interestingly seem to be the result of the uniformly strong repression of a cluster of 21 genes (RS08795–08895) linked to bacterial microcompartments (BMCs) (**Figure S1**). Despite the 4.8-fold (q < 0.01) upregulation of one of the bifunctional aldehyde/alcohol dehydrogenases (RS18395; AdhE1) which catalyses ethanol production directly from acetyl-CoA, ethanol was still most likely produced through acetate using the AOR activity as AOR1 transcripts were ~160-fold more abundant (RPKM) compared to AdhE1 (**Tables S2, S3**) across all experiments and AOR2 (RS00490) was up-regulated 2.5-fold (q < 0.01) (**Tables S4, S5**). This is consistent with metabolic flux data (**Figure 3**) and previous –omics and knockout experiments in acetogens (see above).

**Figure 6.**
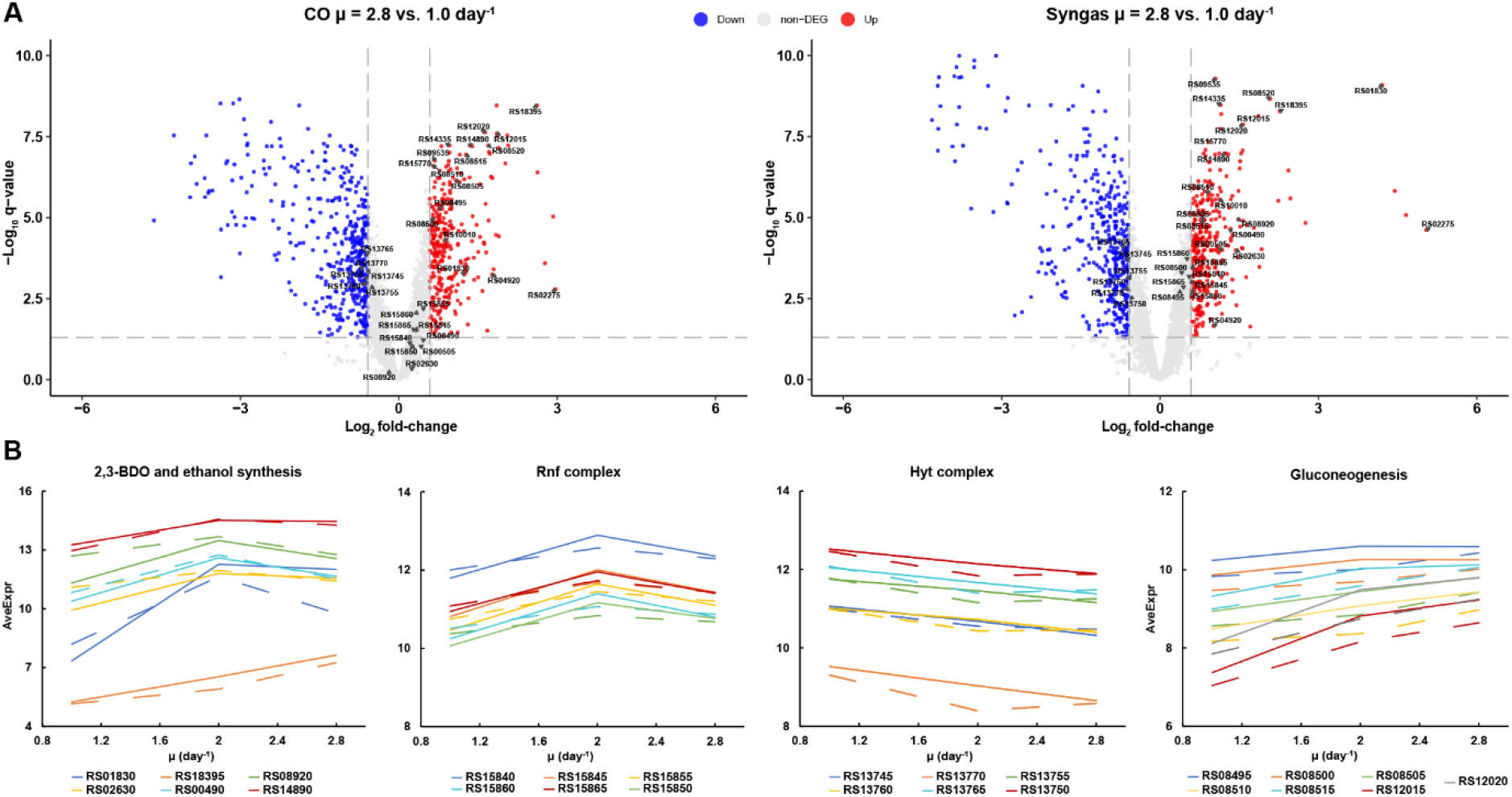
Individual gene expression changes in CO- and syngas-grown *C. autoethanogenum* chemostats. **(A)** Volcano plots showing up-(red) and down-regulated (blue) transcripts for μ = 2.8 vs. 1.0 day^-1^. Genes in **(B)** are indicated by gene ID. Grey lines denote DEG thresholds (fold-change > 1.5 with q-value < 0.05) **(B)** μ-dependent expression profiles for DEGs linked to activities specifically relevant to acetogens (CO, dashed; syngas, solid). *DEG*, differentially expressed gene (fold-change > 1.5 with q-value < 0.05); *μ*, specific growth rate; *AveExp*, average of bioreplicates log_2_ counts per million mapped reads (CPM; see Ritchie et al., 2015). Gene IDs are preceded with CAETHG_. See **Tables S4, S5** for DEG data.

Faster growth demands more energy and this was supported by increased expression (1.5- to 2.2-fold, q < 0.01) of genes of the multi-subunit Fd-NAD^+^ oxidoreductase Rnf complex (RS15845-65) across the studied range of μ (**Figure 6**), as the Rnf complex generates the proton motive force to drive the ATPase in *C. autoethanogenum* (Hess et al., 2016; Tremblay et al., 2012). Notably, up-regulation of NAD(P)- (RS02275) and FAD-dependent (RS04920) oxidoreductases was also detected (7.9- to 32.9-fold and 2.0- to 3.4-fold, respectively, q < 0.01) on both gases that might play a role in the maintenance of redox homeostasis (**Figure S2**). Intriguingly, despite the elevated H_2_ uptake with faster growth on syngas (**Figure 1C**), ~1.5-fold (q < 0.01) repression of the main H_2_ oxidiser – the HytA-E complex (RS13745–70) – (Valgepea et al., 2021; Mock et al., 2015; Wang et al., 2013) components was seen (**Figure 6**). Moreover, similar down-regulation was observed in CO cultures, potentially indicating μ-dependent regulation of the HytA-E complex. Repression of HytA-E also means that gene expression levels at μ = 1.0 day^-1^ were sufficient to realise higher H_2_ uptake with faster growth, which was also the case for higher CO fixation as WLP genes were not detected as DEGs (**Tables S4, S5**). In contrast, increased flux throughput for biomass synthesis with faster growth was supported by elevated expression of gluconeogenesis genes (RS08495–08515, RS12015, and RS12020) (**Figure 6B**). In the future, further work using genetically modified strains is needed to determine whether any of the putative transcription factors that were up-regulated with faster growth (**Figure S2**) are responsible for increased gene expression in any of the cases above.

## DISCUSSION

Exploring the physiological boundaries of acetogens has been informative for understanding regulation of their energy-limited metabolism (Valgepea et al., 2017a; Mahamkali et al., 2020; Jin et al., 2021; Klask et al., 2020). Additionally, various bioprocess approaches, multi-omics analysis, construction of metabolic models, and development of genetic tools are being pursued for development of cell factories with enhanced substrate conversion, product distribution, and expanded product spectrum (Valgepea et al., 2018; Lemgruber et al., 2019; Heffernan et al., 2020; Smith et al., 2020; Fackler et al., 2021; Jin et al., 2021; Bourgade et al., 2021; Diender et al., 2021; Molitor et al., 2019; Pavan et al., 2022). Notably, the effects of μ on acetogen metabolism and the gas fermentation bioprocess have not been established. Here, we investigated the μ-dependent bioprocess performance of the acetogen *C. autoethanogenum* in CO- and syngas-grown steady-state chemostat cultures and assessed metabolic responses using gas analysis, metabolomics, transcriptomics, and metabolic modelling.

We observed higher carbon flux towards ethanol and 2,3-BDO production with faster growth on syngas (**Figure 2**). Increased carbon flux to the same reduced products is also seen with higher steady-state biomass concentrations for syngas-fermenting *C. autoethanogenum* cultures (Valgepea et al., 2017a). Furthermore, we also measured significantly increased specific productivities of ethanol (q_EtOH_) and 2,3-BDO (q2,3-BDO) with faster growth (**Figure 1B**). These trends are beneficial for an industrial gas fermentation process as faster growth and higher biomass levels complimentary lead to higher product yields and volumetric productivities (g/L/h). The fact that no clear trend between μ and carbon flux to 2,3-BDO was seen for CO-grown cultures (**Figure 2**) confirms that feed gas composition has a substantial effect on acetogen product distribution (Valgepea et al., 2018; Jack et al., 2019; Heffernan et al., 2020; Xu et al., 2017; Diender et al., 2016). While cells used the majority of the H_2_ supplied within syngas to directly reduce CO_2_ to formate in the WLP (**Figure 3**), H_2_ supply also had a direct effect on intracellular redox homeostasis as H_2_ oxidation provided up to 10% of the total Fd_re_d generated by the cells (**Table S8**), which is consistent with previous observations in *C. autoethanogenum* (Valgepea et al., 2017a, 2018).

Our transcriptomics data revealed differential expression for more than a thousand genes (DEGs) with increasing μ (**Tables S4, S5**). This means that the acetogen *C. autoethanogenum* uses transcriptional regulation to a large extent at least for facilitating faster growth, compared to previous omics studies that suggest a limited role for hierarchical regulation of gene expression in autotrophic metabolism (Richter et al., 2015; Valgepea et al., 2017a, 2018; Al-Bassam et al., 2018; Mahamkali et al., 2020). In addition to DEGs linked to activities specifically relevant to acetogens (e.g. oxidoreductases, hydrogenases, alcohol/aldehyde dehydrogenases), the functional classification analysis identified genes related to translation within DEGs up-regulated with faster growth (**Figure 5**). This indicates that the well-known positive correlation between μ and total RNA content of biomass (Shaechter et al., 1958) is not sufficient for ensuring faster growth of acetogens either, similarly to μ-dependent datasets for other microbes (Regenberg et al., 2006; Shii et al., 2007; Valgepea et al., 2010, Lahtvee et al., 2011). Notably, COG classification of DEGs identified most genes within the group “Function unknown” (S) (**Figure 5A**), while DEGs also included many genes with unclear functions for autotrophic growth, e.g. sugar, amino acid, and other transporters, hypothetical proteins, BMC-related genes (**Tables S4, S5**). This highlights the need for mapping genotype-phenotype links and improving gene annotations for advancing understanding of acetogen metabolism and engineering of cell factories.

Though our chemostat cultures were CO-limited and thus the residual CO concentration in the liquid phase was low, elevated CO feeding could still inhibit cellular hydrogenases and hence H_2_ uptake for syngas cultures (Thauer et al., 1974; Wang et al., 2013; Mahamkali et al., 2020). Higher syngas, and thus also CO, feeding rates to support faster growth did not, however, prevent elevated H_2_ uptake (**Figure 1C**). Interestingly, expression of genes responsible for realising higher H_2_ uptake – HytA-E for H_2_ oxidation and its complex partner FdhA for direct CO_2_ reduction to formate using H_2_ – was not increased at the same time (**Figure 6**). This is consistent with proteomics data showing no expression change of hydrogenase proteins even when *C. autoethanogenum* increases its H_2_ uptake from near-zero to ~30 mmol/gDCW/h (Valgepea et al., 2018). Similarly, we could neither detect up-regulation of any WLP genes with substantially increased flux through the pathway with faster growth. However, the expression of gluconeogenesis genes was increased to support faster biomass synthesis (**Figure 6**). These observations suggest evolutionary prioritisation of pathways in *C. autoethanogenum* to ensure sufficient “baseline” enzymatic capacity for increasing flux throughput through key catabolic pathways for CO and H_2_ metabolism, consistent with integrative analysis in *C. autoethanogenum* (Valgepea et al., 2021) and other microbes (Daran-Lapujade et al., 2007; Valgepea et al., 2013; Adamberg et al., 2012). This would also provide acetogens an advantage in their natural environments as cells could rapidly respond to CO and H_2_ availability without delays to kick-start metabolism.

Our transcriptomics data are also informative regarding by-product synthesis. In a closely related acetogen – *C. ljungdahlii* – gene expression of all 2,3-BDO production pathway genes is increased prior to 2,3-BDO production in autotrophic batch cultures (Köpke et al., 2011). We, however, detected strong up-regulation (18.5-fold) of only BDH (RS01830; reduces acetoin to 2,3-BDO) with substantially increased carbon flux to 2,3-BDO with faster growth on syngas despite the 2,3-BDO production pathway genes being nearly identical between the two species (Brown et al., 2014). This potentially suggests that there is no transcriptional limitation for channelling carbon from pyruvate to acetoin and makes BDH a protein and metabolic engineering target for improving 2,3-BDO production in *C. autoethanogenum*. While our transcriptomics and metabolic modelling results confirmed the dominant role of AOR activities for ethanol biosynthesis in acetogens (Marcellin et al., 2016; Valgepea et al., 2018; Heffernan et al., 2020; Liew et al., 2017; Richter et al., 2016), strong repression of a cluster of 21 genes (RS08795–08895) linked to BMCs including two putative acetaldehyde dehydrogenases (RS08810 and RS08865) was detected (**Figure S1**). Clustering of putative alcohol and acetaldehyde dehydrogenases within BMC-encoding genes in *C. autoethanogenum* has been noted before (Mock et al., 2015). Cells use BMCs to optimise metabolic pathways by encapsulating enzymes within a protein shell to trap toxic intermediates (Chowdhury et al., 2014; Kerfeld and Erbilgin, 2015). While BMCs seem to play an important role for heterotrophic growth of the acetogen *Acetobacterium woodii* (Schuchmann et al., 2015; Chowdhury et al., 2020; Chowdhury et al., 2021), it remains to be seen how relevant are BMCs for *C. autoethanogenum* autotrophy as the 21 gene cluster was expressed at very low abundances (**Tables S2, S3**).

Overall, our study provides important quantitative information and systems-level analysis of the effects of the key physiological parameter – μ – on acetogen metabolism and the gas fermentation bioprocess during steady-state cultures. We conclude that the bioprocess benefits from faster growth of *C. autoethanogenum* by supporting both higher product yields and productivities. Furthermore, our work advances understanding of transcriptional regulation in acetogens and supports the concept that cells maintain sufficient “baseline” expression of key catabolic pathways for increasing flux throughput. Finally, differential expression of genes with unclear functions emphasises the need for mapping genotype-phenotype links and improving gene annotations for advancing understanding of acetogen metabolism and engineering of cell factories.

## CONFLICT OF INTEREST STATEMENT

LanzaTech has interest in commercial gas fermentation with *C. autoethanogenum*. AH and MK are employees of LanzaTech.

## AUTHOR CONTRIBUTIONS

LAL: Conceptualization, Methodology, Formal analysis, Investigation, Writing – Original Draft, Writing – Review & Editing; HI: Methodology, Formal analysis, Investigation, Writing – Review & Editing; KB: Investigation; KR: Methodology, Formal analysis, Writing – Review & Editing; CB: Software, Resources, Writing – Review & Editing; AH: Resources, Writing – Review & Editing, Project Administration; EM: Resources, Writing – Review & Editing; MK: Conceptualisation, Resources, Writing–Review & Editing; KV: Conceptualization, Methodology, Formal analysis, Investigation, Resources, Writing – Original Draft, Writing – Review & Editing; Supervision, Project Administration, Funding Acquisition.

## FUNDING

This work was funded by the European Union’s Horizon 2020 research and innovation programme under grant agreement N810755 and the Estonian Research Council’s grant agreement PSG289. Australian Government funding through its investment agency, the Australian Research Council, towards the ARC Centre of Excellence in Synthetic Biology (CE200100029) is gratefully acknowledged.

## ACKNOWLEDGMENTS

We thank the following investors in LanzaTech’s technology: BASF, CICC Growth Capital Fund I, CITIC Capital, Indian Oil Company, K1W1, Khosla Ventures, the Malaysian Life Sciences, Capital Fund, L. P., Mitsui, the New Zealand Superannuation Fund, Novo Holdings A/S, Petronas Technology Ventures, Primetals, Qiming Venture Partners, Softbank China, and Suncor.

## SUPPLEMENTARY MATERIAL

The Supplementary Material for this article can be found online at:

## DATA AVAILABILITY STATEMENT

RNA sequencing data have been deposited in the NCBI Gene Expression Omnibus repository under accession number GSEXXX.

