## Supplementary material for "Faster growth enhances low carbon fuel and chemical production through gas fermentation": Figure S1

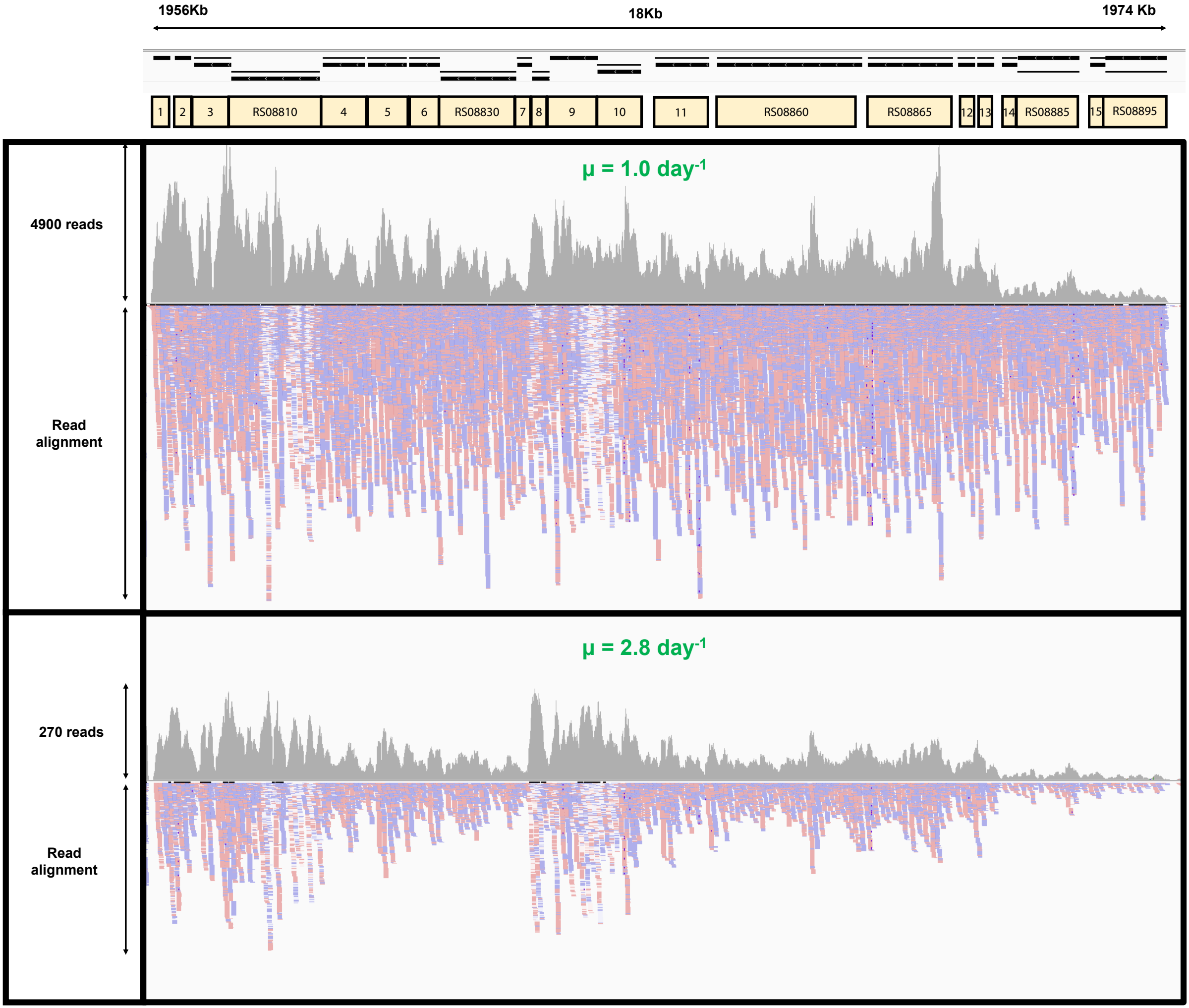


**Figure S1** Faster growth leads to strong repression of a cluster of 21 genes (RS08795–08895) linked to bacterial microcompartments (BMCs). Grey upper part shows the wiggle plot of the sequence read coverage. Blue-pink lower part shows mapping of aligned reads where pink indicates (+) and blue (-) strands. Numbers inside yellow boxes above wiggle plot denote gene IDs: 1–3 (RS08795–RS08805), 4–6 (RS08815–RS08825), 7–11 (RS08825–RS08855), 12–14 (RS08870  RS0880), and 15 (RS08890). *μ*, specific growth rate. Gene IDs are preceded with CAETHG_.
