## Supplementary material for "Faster growth enhances low carbon fuel and chemical production through gas fermentation": Figure S2

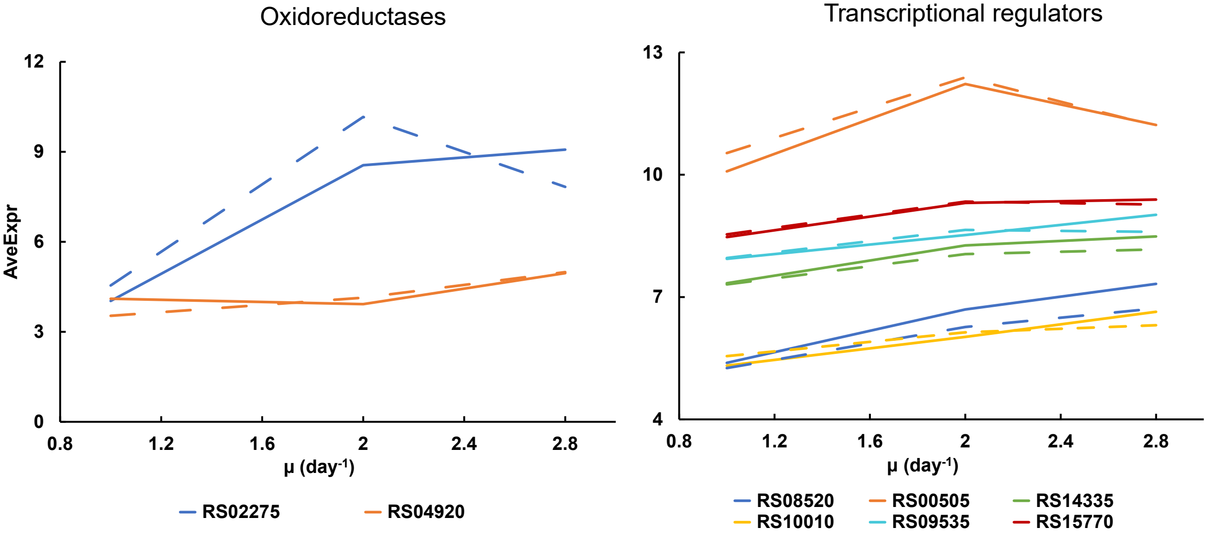


**Figure S2** Individual gene expression changes in CO- and syngas-grown *C. autoethanogenum* chemostats. μ-dependent expression profiles for DEGs of oxidoreductases and transcriptional regulators (CO, dashed; syngas, solid). *DEG*, differentially expressed gene (fold-change > 1.5 with q-value < 0.05); *μ*, specific growth rate; *AveExp*, average of bioreplicates $\log_2$ counts per million mapped reads (CPM; see Ritchie *et al*., 2015). Gene IDs are preceded with CAETHG_. See Tables S4, S5 for DEG data.
